# The wild-type hERG cryo-EM structure fails to support conduction and evolve toward an inactivated-like selectivity filter conformation in molecular dynamics simulations

**DOI:** 10.64898/2026.09.02.748792

**Authors:** Maria Vittoria Leonardi, Antonio Michelucci, Yessenbek K. Aldakul, Han Sun, Luigi Catacuzzeno, Simone Furini

**Affiliations:** Department of Chemistry, Biology and Biotechnology, University of Perugia, Italy; Leibniz-Forschungsinstitut für Molekulare Pharmakologie (FMP) & Department of Chemistry, Technische Universität Berlin, Berlin, Germany; Department of Electrical, Electronic and Information Engineering "Guglielmo Marconi”, University of Bologna, via dell’Università 50, Cesena (FC), 47521, Italy

## Abstract

The atomic structure of the human Ether-à-go-go–Related Gene (hERG) K^+^ channel has recently been resolved by cryo-electron microscopy (cryo-EM) under both high- and low-K^+^ conditions, in order to obtain information on the mechanism of the K^+^-sensitive, very rapid C-type inactivation typical of this channel. Although the currently available high-K^+^ structures have been widely interpreted as representing the conductive, active state, whether they correspond to a dynamically stable conductive conformation remains unresolved. Here, we used extensive all-atom molecular dynamics (MD) simulations, with and without Electronic Continuum Correction (ECC), to investigate selectivity filter (SF) dynamics and ion permeation in wild-type (WT) hERG and the non-inactivating N629D mutant. Across all membrane potentials tested, WT hERG failed to support K^+^ permeation and instead spontaneously evolved towards a non-conductive SF conformation characterized by extracellular dilation, localized inner constriction, depletion of the outer ion-binding sites, and persistent trapping of K^+^ ions within the central binding sites of the filter, closely resembling the inactivated SF of Shaker channels recently resolved by cryo-EM. By contrast, N629D maintained a stable conductive SF architecture, analogous to the conductive filters of canonical K^+^ channels such as KcsA, while exhibiting robust voltage-dependent K^+^ permeation. ECC enhanced ion permeation in N629D mutant, but failed to support conduction in WT hERG, which remained structurally and functionally non-conductive. Together, these findings challenge the prevailing interpretation of the high-K^+^ WT cryo-EM structure as a stable conductive state and identify SF remodeling as the structural mechanism underlying hERG C-type inactivation.

## Introduction

The human Ether-à-go-go–Related Gene (hERG) channel is a voltage-gated K^+^ channel that mediates the rapid delayed rectifier current, a major determinant of cardiac action potential repolarization and normal cardiac rhythm. A distinctive feature of hERG is its unusually rapid C-type inactivation, which develops faster than channel activation and strongly limits outward K^+^ current during depolarization. Upon membrane repolarization, hERG channels recover rapidly from inactivation while deactivating more slowly, generating the characteristic tail currents that contribute substantially to the final phase of cardiac repolarization (Vandenberg et al. 2012). This distinctive gating behavior has made hERG a paradigmatic system for investigating the interplay between ion permeation and selectivity filter (SF) gating.

A large body of evidence from hERG and other K^+^ channels indicates that C-type inactivation originates within the SF, the narrow pore region responsible for ion selectivity and conduction. Ion permeation and inactivation are tightly coupled processes: changes in ion occupancy influence SF stability, while conformational rearrangements directly regulate conduction. Understanding how these processes are coupled is therefore essential for elucidating the molecular basis of hERG gating.

The structural basis of C-type inactivation has been extensively investigated in several K^+^ channels and is now described by two principal mechanistic paradigms. In the bacterial channel KcsA, the inactivation is associated with collapse of the SF, involving progressive backbone rearrangements that constrict the permeation pathway, disrupt the central ion-binding sites through carbonyl reorientation, and ultimately abolish K^+^ conduction (Zhou et ak., 2001; Cuello et al. 2010). By contrast, studies of the eukaryotic Shaker K^+^ channel have shown that C-type inactivation occurs without collapse of the SF. Early electrophysiological experiments demonstrated that removal of extracellular K^+^ unmasks residual Na^+^ permeation through the inactivated state of the constitutive inactive W434F mutant (Starkus et al. 1998), a finding incompatible with an occluded SF. More recently, atomic-resolution cryo-electron microscopy (cryo-EM) structures suggested that inactivation is associated with extracellular dilation of the SF rather than a collapse (Tan et al. 2022; Stix et al. 2023). Together, these studies demonstrate that structurally distinct SF conformations can produce the same functional outcome—loss of K⁺ conduction.

The structural mechanism underlying hERG inactivation remains even more unclear. Functional studies indicate that hERG inactivation involves multiple conformational states rather than a single structural transition. In the complete absence of K^+^, inactivated hERG channels transiently conduct Na^+^ before slowly evolving toward a more stable non-conductive state (Gang e Zhang 2006), indicating that loss of K^+^ conduction precedes complete pore inactivation. Such behavior is difficult to reconcile with a simple binary transition between conductive and collapsed filters and instead supports a model in which distinct SF conformations exhibit different permeation properties.

Recent cryo-EM studies have substantially advanced the structural understanding of hERG gating. The first high-resolution structure, determined under high K^+^, revealed a SF with the canonical cylindrical geometry and S0-S4 binding sites characteristic of conductive K^+^ channels, and has therefore been widely interpreted as representing the conductive state (Wang e MacKinnon 2017; Lau et al. 2024). In contrast, a structure resolved under low-K^+^ conditions displays pronounced rearrangements within the SF, including reorientation of the V625 carbonyl groups away from the pore axis, disruption of the central ion-binding sites, and stabilization of a non-conductive conformation strongly influenced by interactions involving S620 (Lau et al., 2024). Because extracellular K^+^ stabilizes the conductive state of hERG, the low-K^+^ structure has been proposed to represent an inactivated conformation. However, whether the high-K^+^ structure corresponds to a dynamically stable conductive state or instead occupies an early intermediate along the inactivation pathway remains unresolved.

Recent molecular dynamics (MD) simulations of hERG channel performed under equilibrium conditions revealed that the WT SF can spontaneously rearrange toward dilated conformations, closely resembling the inactivated state of Shaker channels (Pettini et al. 2023). This raises the possibility that the high-K^+^ cryo-EM structure of hERG channel is intrinsically predisposed towards inactivation, and a direct computational assessment of whether this structure can sustain K^+^ conduction under experimentally relevant membrane potentials is still lacking. Previous MD studies reporting K^+^ permeation through hERG generally relied either on restraints applied directly to the SF backbone or on extremely large membrane potentials (i.e., ±700 mV) (Miranda et al. 2020; Lau et al. 2024; Ngo et al. 2025), both of which can perturb the intrinsic dynamics of the SF. Moreover, studies of Shaker have demonstrated that sufficiently strong electric fields can drive ion permeation even through structurally inactivated SF conformations (Stix et al. 2023), complicating interpretation of conduction under non-physiological conditions. Because ion occupancy within the SF is strongly voltage-dependent and tightly coupled to C-type inactivation, determining whether the WT hERG structure sustains K^+^ conduction under moderate membrane potential is therefore essential for establishing its physiological relevance.

Here, we investigated the ion conduction properties and SF dynamics of the WT hERG and the non-inactivating N629D mutant using extensive, unrestrained all-atom MD simulations with different force field parameters under membrane potentials close to those employed experimentally. By combining analyses of ion permeation, ion occupancy, and SF conformational dynamics, we examined whether the currently available high-K^+^ WT hERG cryo-EM structure represents a stable conductive state or instead spontaneously transitions towards a non-conductive, inactivated-like SF conformation. Together, these simulations provide a framework to directly link SF architecture and dynamics to the functional mechanisms underlying hERG conduction and inactivation.

## Methods

### Atomic models

MD simulations were performed starting with the WT HERG high-K^+^ structure pdb: 9CHP (Lau et al. 2024). Only the pore region of the channel was included in the model, from residue Tyr545 to residue Tyr667 and ACE and NME capping were applied respectively at N-terminal and C-terminal regions. The model of hERG-N629D was built using the option in CHARMM-GUI “mutate residue”. The system was embedded in a membrane patch of 158 POPC (1-palmitoyl-2-oleoyl-sn-glycero-3-phosphocholine) using CHARMM-GUI (Jo, S., et all, 2008). The system was solvated using TIP3P water molecules model, and 600 mM of KCl were added. Potassium ions were manually placed at binding sites S4, S3, S2, S1 in the SF.

### MD simulations

All simulations used the Amber14SB force field (Maier et al. 2015). Van der Waals interactions were truncated at 9 Å. Standard AMBER scaling of 1-4 interactions was applied. Long-range electrostatic interactions were calculated with the Particle Mesh Ewald method using a grid spacing of 1.0 Å (Essmann et al. 1995). The SETTLE algorithm was used to restrain bonds with hydrogen atoms (Tuckerman et al. 1992). Electronic Continuum Correction (ECC) was applied to all the ions and charged residues in a set of simulations using the Amber14SB force field following the exact method reported in the work from De Groot for each force field (Hui et al. 2024). The charge scaled parameters for ions and residues were downloaded from https://github.com/deGrootLab/Charge_Scaling_in_Potassium_Channel_Simulations_paper and replaced in the standard force field. The temperature was controlled at 310 K by coupling to a Langevin thermostat with a damping coefficient of 1 ps^−1^. A pressure of 1 atm was maintained by coupling the system to a Nose−Hoover Langevin piston, with a damping constant of 25 ps and a period of 50 ps (Feller et al. 1995). GROMACS was used for all the simulations (Abraham et al. 2015). The equilibration protocol consisted of 5.000 steps of energy minimization, followed by 10 ns of dynamics in the NPT ensemble with a 2 fs timestep and position restraints in all the heavy atoms, then a 50 ns in the NPT ensemble with a 2 fs timestep with no position restraints. Subsequently, all the production runs were carried out in 4 replicas from 2 up to 5 μs. Electric field was applied parallel to the z-axis of systems to set a membrane potential of ±200mV and ±400mV.

### Analysis of the MD trajectories

Trajectory analyses were performed with custom Python code using MDanalysis (Michaud-Agrawal et al. 2011) and the NumPy and SciPy ecosystem. Ions and water molecules were considered in binding sites S0-S4 when the axis coordinate was in between the center of the two layers of oxygen atoms delimiting the corresponding site. Ramachandran angles of backbone amino acids were used to evaluate the distortion of SF with respect to the cryo-EM structure. An analysis was performed with RMSD and RMSF to assess structural flexibility and the average distance of opposing carbonyl oxygen atoms, calculated as a mean distance during the simulations, were used to assess the distortion of SF sites. VMD was used to inspect trajectories and to generate images of the systems (Humphrey et al. 1996).

## Results

### The high-K^+^ WT hERG cryo-EM structure fails to support K^+^ conduction under moderate membrane potentials

To assess whether the experimentally resolved high-K^+^ hERG structure supports ion conduction, we performed extensive all-atom MD simulations starting from the cryo-EM structure reported by Lau and colleagues (PDB ID: 9CHP). Simulations were carried out at applied membrane potentials of ±200 and ±400 mV using the Amber14SB force field. Since ion occupancy is not resolved in the cryo-EM structure, all simulations were initiated from the canonical four-ion configuration with K^+^ ions occupying the S1-S4 binding sites, a configuration supported by crystallographic ion density maps and previous computational studies of other K^+^ channels (Lee et al. 2025; Köpfer et al. 2014).

Across all simulations, the WT hERG channel consistently failed to support K^+^ permeation. Shortly after equilibration and release of positional restraints, two K^+^ ions vacated the extracellular S0 and S1 binding sites, after which the remaining two K^+^ ions remained stably trapped within the central S2-S3 region throughout the trajectories (Figure 1). No complete permeation event was observed under any membrane potential, including high transmembrane voltages of ±400 mV (Table S1). A single trajectory obtained at -400 mV exhibited limited ion rearrangement, in which one extracellular K⁺ ion reached the S2 site, while another occupied the S4-S3 boundary (Figure 1). However, this configuration is rarely observed with productive permeation in conductive K^+^ channels (Domene et al. 2021a; Hui et al. 2024) and did not progress to complete translocation across the filter. We therefore do not consider it a *bona fide* permeation event. The absence of conduction is difficult to reconcile with experimentally measured single-channel conductance of hERG, (2-10 pS) (Kiehn et al. 1996; Vijayvergiya et al. 2015), which corresponds to 50-250 permeation events over the cumulative simulation time of 20 μs at 200 mV. Even accounting for the well-established tendency of classical additive force fields to underestimate the K^+^ conductance by approximately one order of magnitude (Hui et al. 2024), several events would still be expected at ±200 (5-25) mV and at ±400 mV (10-50). The complete absence of such events therefore strongly indicates that the simulated WT cryo-EM structure does not represent a stable conductive state.

**Figure 1.**
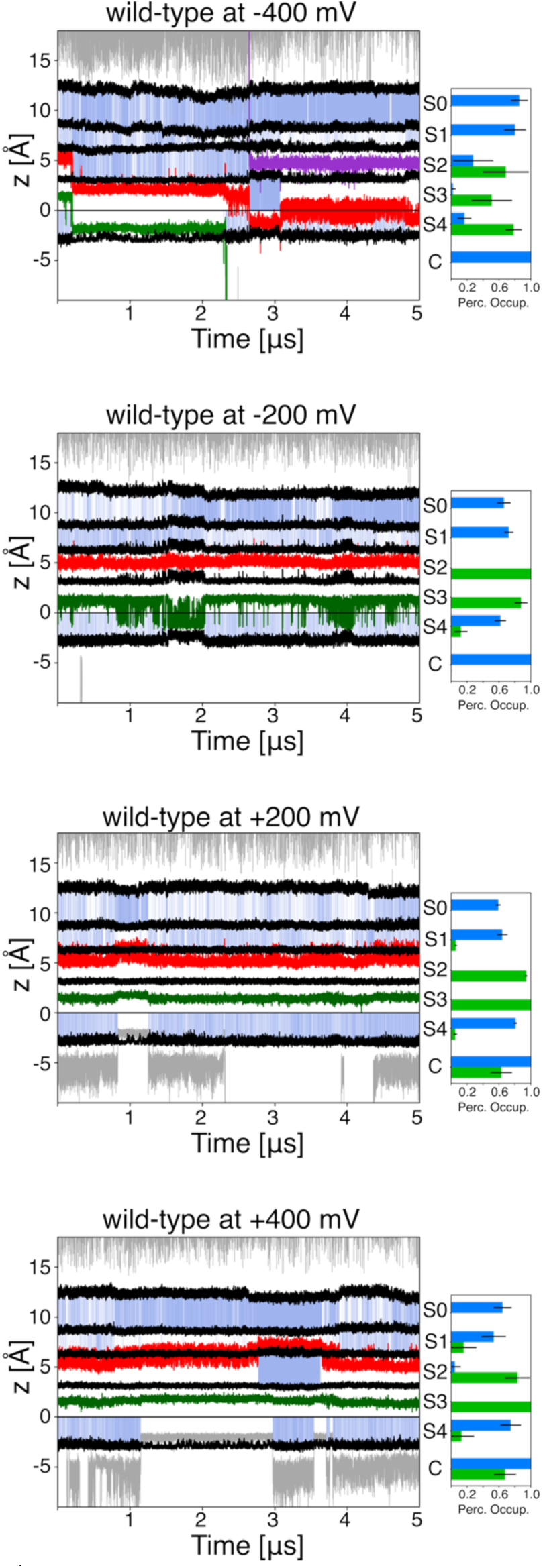
Representative K^+^ trajectories and site occupancies in WT hERG. (Left panels) The position of K^+^ ions along the pore axis is shown for one representative replica among the four simulations performed at each membrane potential. Grey lines indicate ions outside the SF, whereas colored lines indicate ions occupying the SF. Blue shading indicates S0-S4 binding sites, which are occupied by water molecules. Black lines indicate the boundaries between adjacent SF binding sites. (Right panels) The percentage occupancy of the cavity (C), and binding sites in the SF (S4-S0), is reported for potassium ions (green) and water molecules (blue). Error bars represent the standard deviation calculated across the four independent replicas. The simulations were performed with standard Amber14SB force field.

Average ion occupancy profiles further support this conclusion (Figure 1). The SF remained predominantly occupied at the central S2 and S3 sites, whereas occupancy of the extracellular S0 and S1 sites was markedly reduced. Consistent with this depletion, water molecules frequently populated the outer region of the SF and replaced K^+^ ions at the S0-S1 region, disrupting the multi-ion occupancy pattern required for efficient K^+^ knock-on conduction (Köpfer et al. 2014; Mendoza Uriarte e Roux 2026; Domene et al. 2021b).

### The non-inactivating N629D mutation stabilizes a conductive state

To test whether the lack of conduction observed in the WT channel results from spontaneous relaxation of SF toward an inactivated-like filter conformation, we performed an equivalent set of simulations of the N629D mutant with standard Amber14SB force field. This mutation introduces a negatively charged residue immediately above the extracellular mouth of the SF and is well known to abrogate fast inactivation (Lees-Miller et al. 2000).

In sharp contrast to WT channel, the N629D mutant supported K^+^ permeation under the same simulation conditions. At ±200 mV, only partial permeation events were sampled, whereas increasing the membrane potential to ±400 mV produced frequent complete translocation events (Table S1). Representative trajectories at -400 mV revealed continuous exchange of K^+^ ions across the filter, in marked contrast to the persistent trapping of ions within the S2-S3 region observed in the WT hERG (Figure 2).

**Figure 2.**
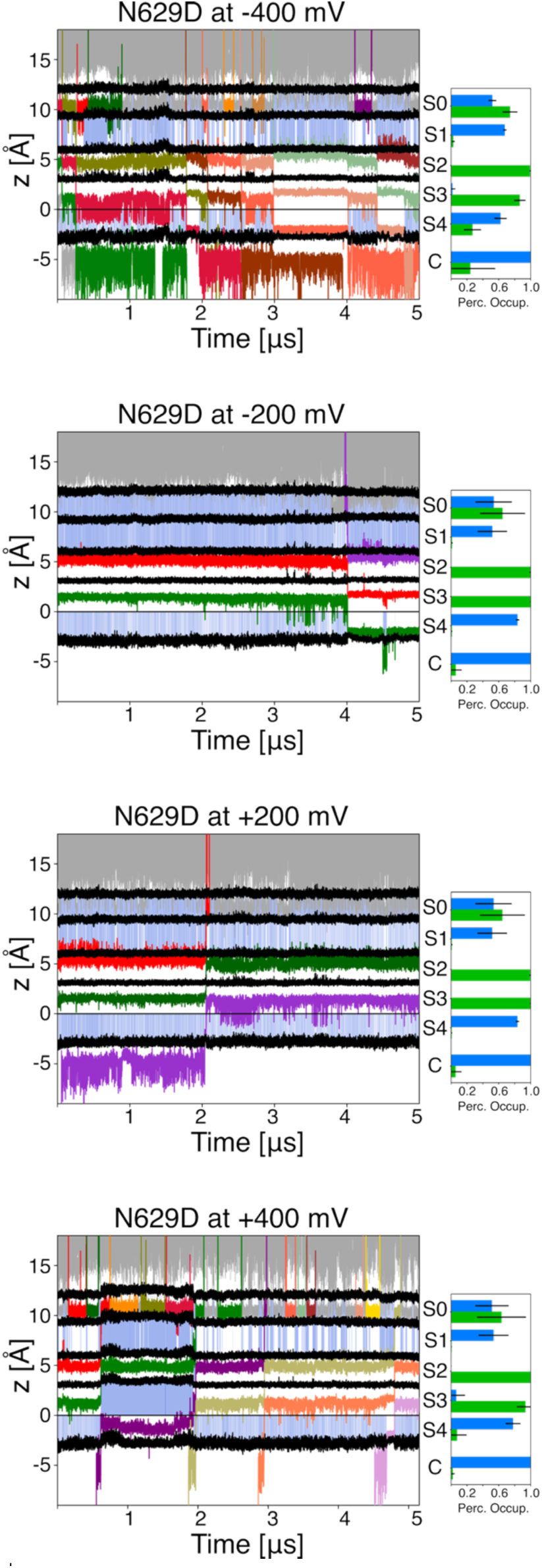
Representative K^+^ trajectories and site occupancies in N629D hERG. (Left panels) The position of K^+^ ions along the pore axis is shown for one representative replica among the four simulations at each membrane potential. Grey lines indicate ions outside the SF, whereas colored lines indicate ions occupancy inside the SF. Blue shading indicates the S0-S4 binding sites, which are occupied by water molecules. Black lines indicate the boundaries between binding sites of the SF. (Right panels) The percentage occupancy of the cavity (C), and binding sites in the SF (S4-S0), is reported for potassium ions (green) and water molecules (blue). Error bars represent the standard deviations calculated across the four independent replicas. The simulations were performed with Standard Amber14SB force field.

To identify the structural basis of this recovered permeation, we analyzed ion occupancy throughout the SF. Compared with WT channel, the N629D mutant exhibited significantly higher occupancy of the extracellular portion of the filter, particularly at the S0 binding site. This redistribution is consistent with electrostatic stabilization of K^+^ ions by the negatively charged D629 side chain, promoting the multi-ion occupancy required for efficient knock-on permeation. Water occupancy within the SF was broadly similar in WT and N629D hERG, with water molecules predominantly localized near the S1 and S4 sites. Nevertheless, transient water permeation events accompanying ion translocation were also observed, indicating that conduction in N629D is compatible with partially hydrated permeation states.

Interestingly, K^+^ conduction in N629D was strongly voltage-dependent, with substantially higher permeation rates observed at negative membrane potentials. A possible explanation is the asymmetric availability of ions on the two sides of the SF, which directly influences the efficiency of the knock-on mechanism. The intracellular cavity of hERG is unusually narrow and hydrophobic, properties that may limit intracellular ion availability (Naranjo et al. 2016; Wang e MacKinnon 2017). By contrast, the extracellular vestibule is directly exposed to the bulk solution and, in the N629D mutant, further concentrates K^+^ ions through electrostatic attraction by the D629 side chain. Together, these factors are expected to facilitate extracellular ion loading and enhance inward permeation. Consistent with this interpretation, inward rectification has also been reported experimentally for WT hERG channels (Zou et al. 1997; Kiehn et al. 1996), although comparable single-channel measurements are not yet available for the N629D mutant. Overall, these results demonstrate that suppression of inactivation through the N629D mutation restores stable multi-ion occupancy and robust K^+^ permeation strongly support the conclusion that the non-conductive behavior of the WT hERG arises from spontaneous remodeling of the SF into an inactivated-like conformation.

### Distinct SF architectures underlie the different conductive behavior between WT and N629D mutant hERG

To determine the structural basis of the divergent conductive properties of WT and N629D hERG, we analyzed the SF dynamics throughout the MD trajectories. Whereas the high-K^+^ WT cryo-EM structure rapidly departed from its initial conformation and evolved towards a distinct non-conductive SF architecture, the N629D mutant remained structurally stable, including at lower transmembrane potentials of ±200 mV. Consistent with this behavior, the SF backbone RMSD relative to the cryo-EM structure (PDB ID: 9CHP) was substantially higher in WT than in N629D (Figure 3, filled boxes). Conversely, when the conductive KcsA structure (PDB ID: 1K4C) was used as the reference, the SF backbone RMSD decreased for N629D but increased in WT hERG (Figure 3, empty boxes), indicating that N629D converges toward a canonical conductive filter architecture, whereas WT diverges from it. Residue-level fluctuation analysis further showed that the enhanced mobility of the WT hERG was concentrated at the extracellular mouth of the filter, particularly around residues F627 and G628, whereas this region remained stable in N629D (Figure 3B).

**Figure 3.**
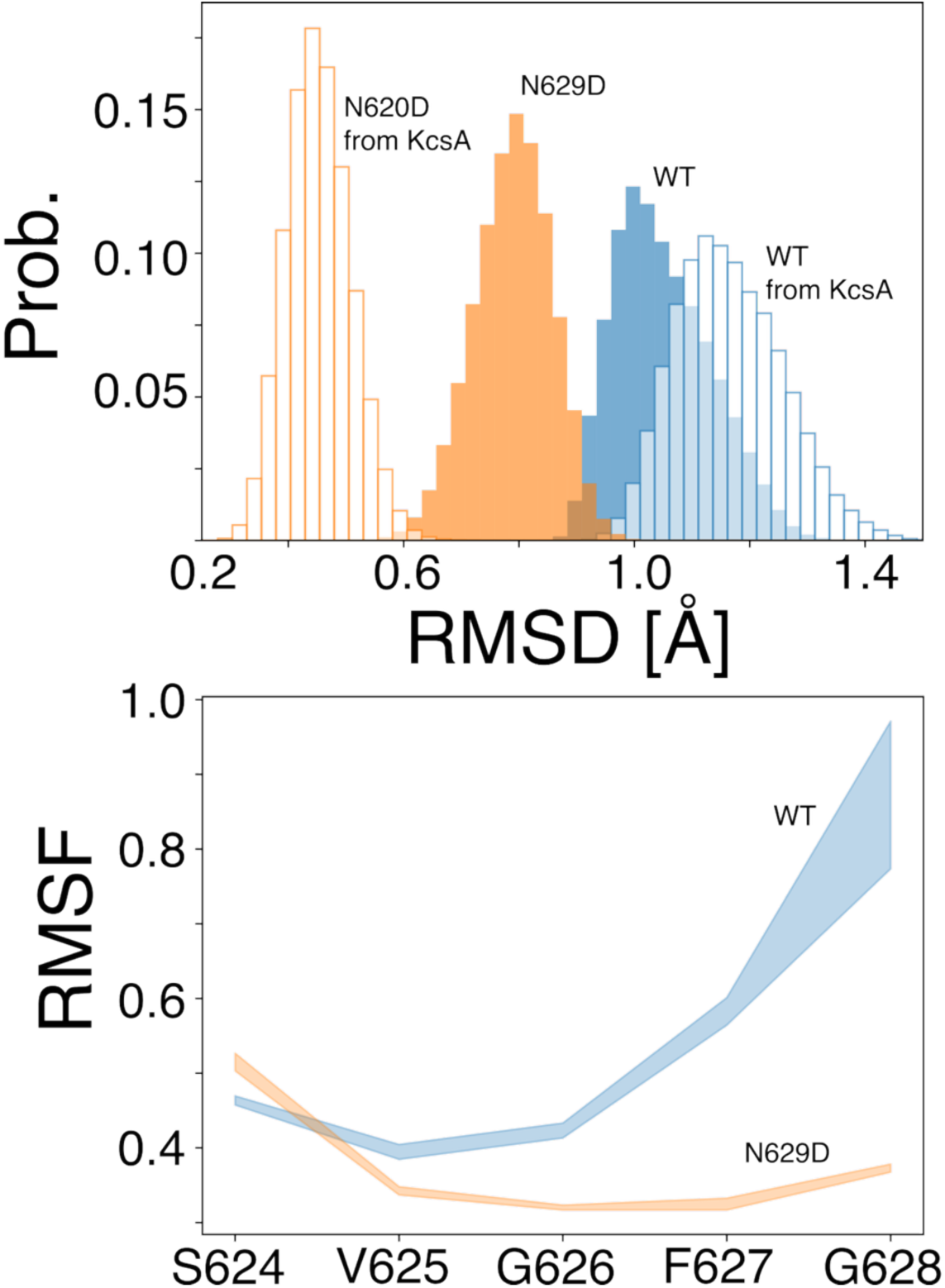
Structural stability and flexibility of the SF in WT and N629D hERG. (Upper panel) Probability distributions of the RMSD of the SF backbone atoms. Filled boxes represent the RMSD relative to cryo-EM structure (PDB ID: 9CHP), whereas empty boxes represent the RMSD relative to the X-ray structure of the KcsA channel in the conductive state (PDB ID: 1K4C). (Lower panel) RMSF of the alpha-carbon atoms of SF residues. Data were calculated from the four independent replicas simulated at +200 mV with Amber14SB force field.

To define the structural determinants underlying this divergence, we analyzed the geometry of the SF along its permeation axis. In WT hERG, the backbone ψ angle of F627 underwent a pronounced reorientation, causing the corresponding carbonyl groups to rotate away from the permeation axis into a tangential orientation. This “carbonyl flipping” disrupts the coordination geometry of the outer ion-binding sites and is consistent with the rapid depletion of K^+^ ions from the S0-S1 region observed in the simulated trajectories. By contrast, F627 remained in its conductive orientation in the N629D mutant, preserving the ion-coordination environment required for permeation (Figure 4).

**Figure 4.**
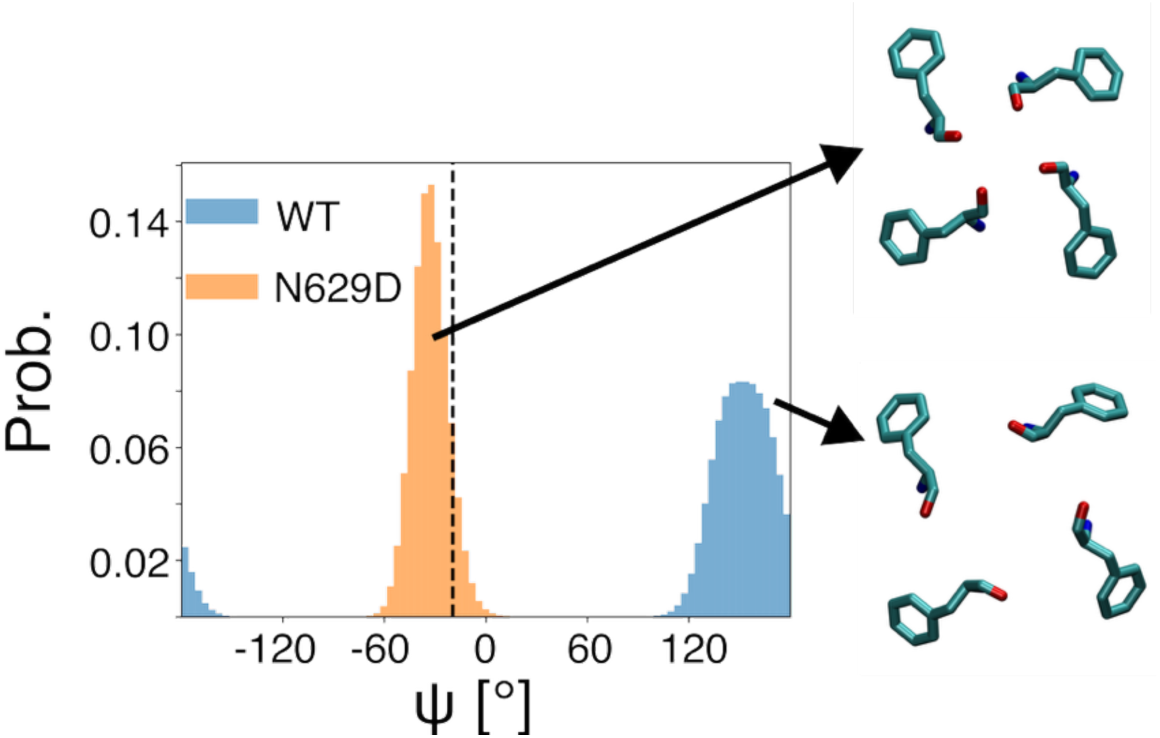
Probability distribution of the backbone ψ angles of residues F627. The dashed horizontal line indicates the value observed in the cryo-EM structure. Representative configurations of the F627 residues, viewed from the extracellular side of the channel, are shown in licorice representation. The F627 carbonyl oxygens are oriented towards the pore axis in the N629D mutant but point laterally in the WT channel. Probability distributions were calculated from the four independent replicas simulated at +200 mV with Amber14SB force field.

Comparison with the conductive/non-inactivating E71A KcsA structure further revealed the structural basis of the different functional states between the two hERG channels. The most prominent difference occurred at the extracellular entrance of the filter. WT hERG developed a marked dilation around F627, due to outward displacement of carbonyl oxygens and loss of S1-S0 coordination, whereas the N629D mutant maintained a stable geometry with inter-subunit oxygen distances of approximately 5 Å, closely matching conductive KcsA (Figure 5).

**Figure 5.**
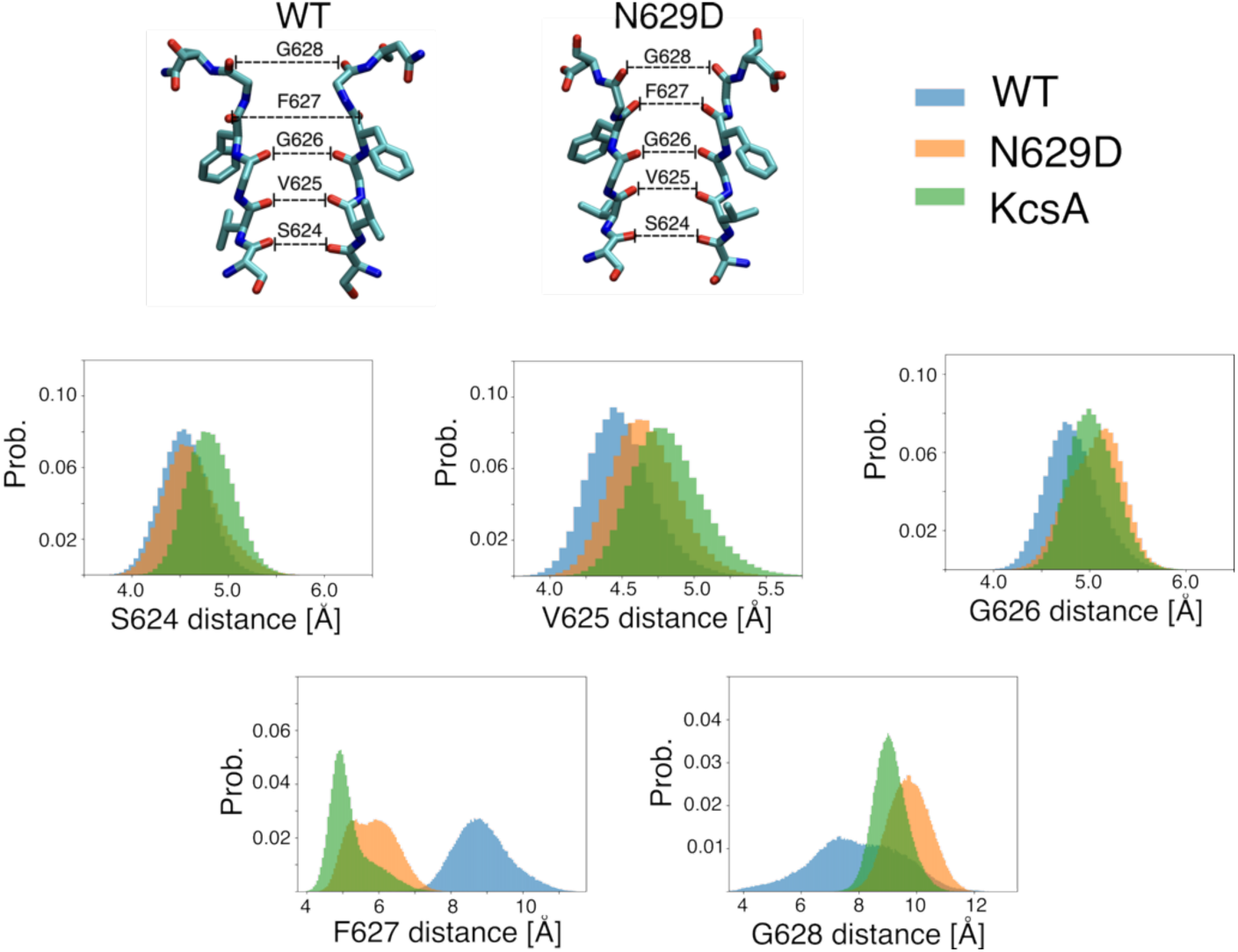
SF architecture of WT and N629D hERG compared with a conductive K^+^ channel (KcsA). Representative structure of the SFs of the WT and N629D hERG channels. Residues 624-629 from two opposing subunits are shown in licorice representation. Probability distributions of the distances between carbonyl oxygen atoms of equivalent residues (S624, V625, G626, F627, and G628) from two opposing subunits are for WT hERG (blue), N629D hERG (orange), and KcsA (green) carrying the E71A mutation. Data were obtained from simulations performed at +200 mV for all channels using Amber14SB force field. KcsA data were taken from (Catacuzzeno et al. 2024) using Amber14SB force field.

A second major divergence was observed near V625 and G626, where WT hERG exhibited a localized constriction of the permeation pathway, reducing the inter-carbonyl distance to approximately 4.5 Å and thereby likely increasing the energetic barrier to efficient K^+^ conduction. This constriction was absent in N629D, which preserved the wider geometry characteristic of conductive K^+^ channels (Figure 5).

Together, these structural rearrangements generate a highly non-uniform SF architecture in WT hERG, combining extracellular dilation with central constriction. By contrast, N629D maintains a canonical conductive filter geometry, providing a structural explanation for its preserved ion occupancy pattern and robust K⁺ permeation.

### Electronic Continuum Correction enhances the functional divergence between WT and N629D hERG channels

To evaluate whether the different behaviors of WT and N629D hERG are dependent on the electrostatic treatment, we repeated the simulations using the Electronic Continuum Correction (ECC) framework (Hui et al., 2025). ECC did not alter the qualitative difference between the two channels but further enhanced their functional divergence. WT hERG remained non-conductive under ECC conditions. In most trajectories, the SF retained the same distorted architecture observed with the standard force field, characterized by persistent F627 carbonyl reorientation, V625 constriction, and trapping K^+^ ions within the central S2-S3 binding sites, as shown in the ion probability distribution in Figure 6 and in Supplementary Material (Figures S1-S3). In three of four simulations performed at -400 mV, the SF instead underwent a more extensive destabilization, characterized by a rapid loss of ion occupancy, water entry, and severe backbone distortions (Figure S4 in Supplementary Material). Although two ions translocation events occurred during these trajectories, they were associated with disruption of the canonical binding sites and highly distorted filter geometries, resembling the inactivated configuration of KcsA (Li et al. 2018). Because these events did not occur through an intact conductive filter, we did not consider them as representative K^+^ permeation.

**Figure 6.**
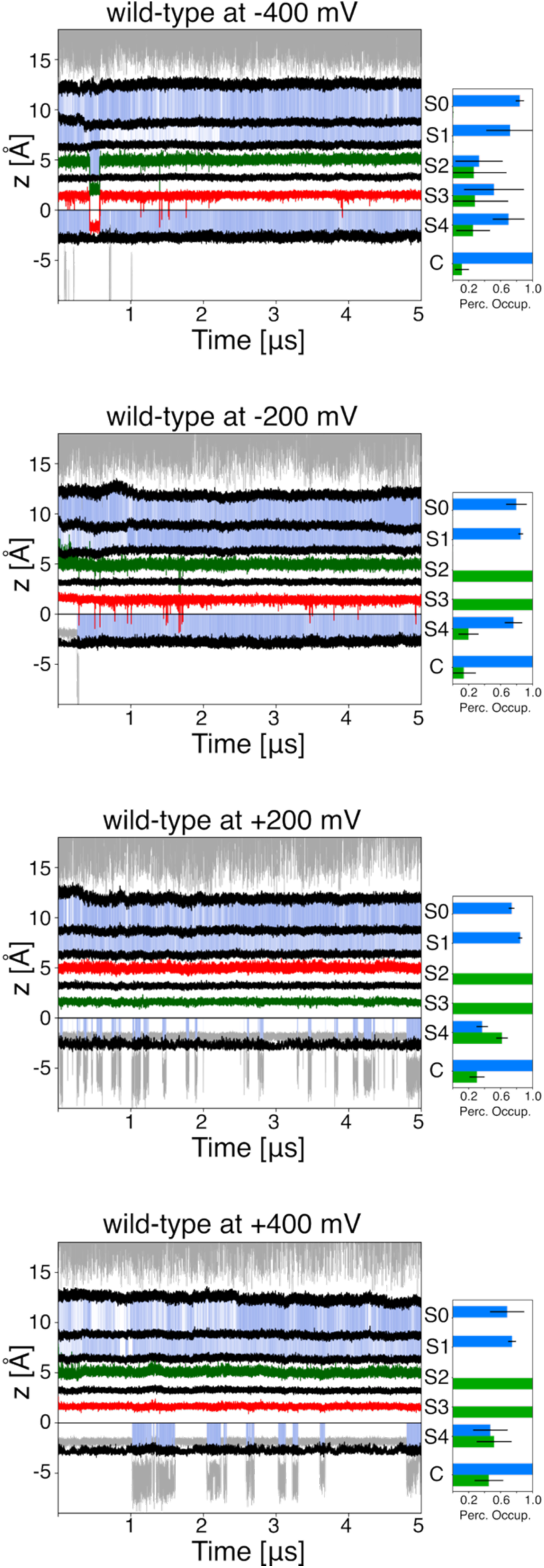
Representative K^+^ trajectories and site occupancies in WT hERG in simulations using Amber14SB with ECC parameters. (Left panels) The position of K^+^ ions along the pore axis is shown for one representative replica among the four simulations at each membrane potential. Grey lines for ions outside the SF, whereas colored lines indicate ions occupying the SF. Blue shading indicates S0-S4 binding sites, which are occupied by water molecules. Black lines indicate the boundaries between adjacent SF binding sites. (Right panels) The percentage occupancy of the cavity (C) and the SF binding sites (S4-S0) is reported for K^+^ ions (green) and water molecules (blue). Error bars represent the standard deviations calculated across the four independent replicas.

On the contrary, ECC substantially enhanced K^+^ permeation in the N629D mutant (3 events per µs with ECC vs. 1.65 events per µs without ECC at -400 mV). Frequent translocation events occurred across a stable conductive SF, indicating that the increased conduction resulted from improved ion dynamics rather than from conformational rearrangement. Moreover, conduction remained strongly voltage-dependent, with higher permeation rates at negative potentials, consistent with the behavior observed using the standard force field (Figure 7).

**Figure 7.**
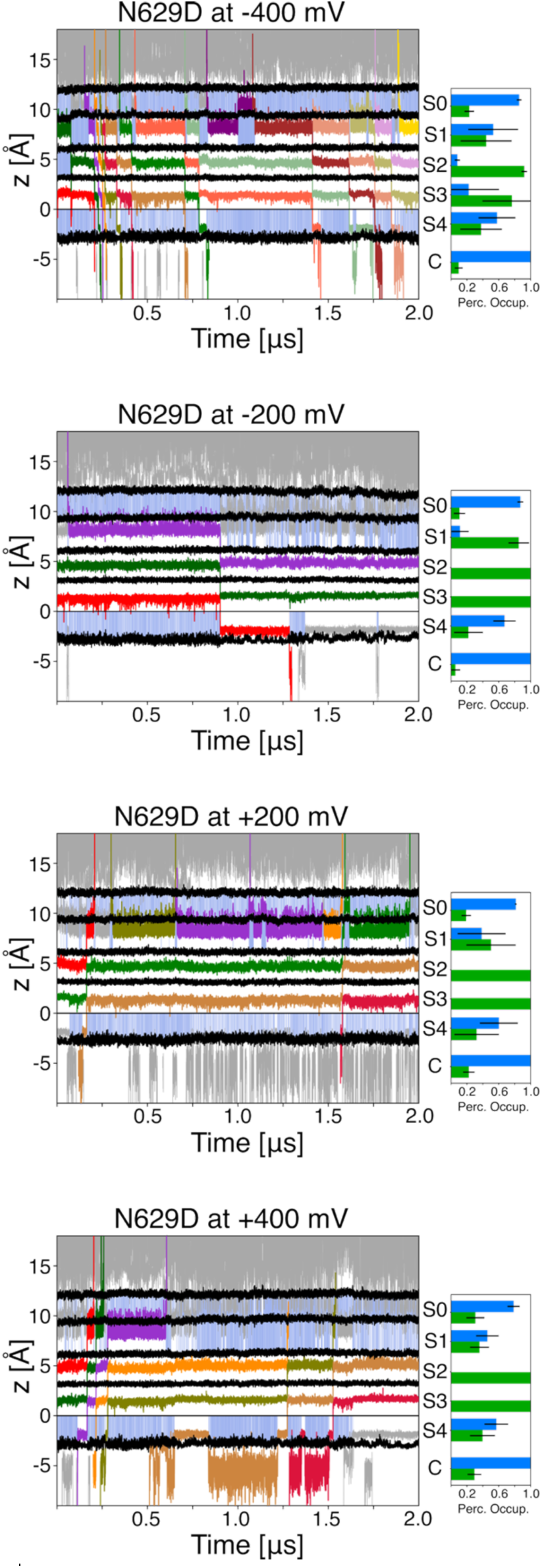
Representative K^+^ trajectories and site occupancies in N629D hERG in simulations with ECC parameters. (Left panels) The position of K^+^ ions along the pore axis of the channel is shown for one of the four simulated replicas at each membrane potential; using grey lines for ions outside the SF, and lines of different colors for ions inside the SF. Blue shading indicates binding sites, S0-S4, occupied by water molecules. Black lines correspond to the boundaries between binding sites of the SF. (Right panels) The percentage occupancy of the cavity (C) and binding sites in the SF (S4-S0), is reported for potassium ions (green) and water molecules (blue). The error bars were calculated as standard deviations among replicas.

Overall, these results demonstrate that ECC preserves and strengthens the functional distinction between WT and N629D hERG. While partial inclusion of electronic polarization enhances permeation in the conductive mutant, it fails to restore conduction in WT, indicating that the lack of permeation in WT hERG arises from its intrinsic non-conductive SF architecture rather than from limitations of the electrostatic model.

## Discussion

### The high-K^+^ WT hERG cryo-EM structure is not a stable conductive state

The central aim of this study was to determine whether the currently available high-K^+^ cryo-EM structure of WT hERG represents a functional conductive conformation capable of supporting K^+^ permeation under moderate, experimentally accessible membrane potentials. Although this structure has been widely interpreted as representing a conductive, active state, its functional identity remained uncertain (Wang and MacKinnon, 2017; Lau et al., 2024). This question is particularly important because hERG channels may possess multiple, functionally different inactivated stats, and experimentally resolved structures may therefore capture only part of its conformational landscape.

Here, extensive all-atoms MD simulations demonstrate that the high-K^+^ WT hERG does not maintain a stable conductive architecture observed experimentally. Instead, the SF rapidly evolves toward a distinct non-conductive conformation characterized by depletion of the outer ion-binding sites, persistent trapping of K^+^ ions within the central S2-S3 region, and the absence of sustained K^+^ permeation. In contrast, the non-inactivating N629D mutant preserves a conductive SF architecture and supports robust voltage-dependent conduction under the same conditions. Together, these findings indicate that the high-K^+^ structure represents a metastable conformation intrinsically prone to the transition towards an inactivated-like, non-conductive state.

### The lack of conduction in WT hERG does not result from insufficient sampling or limitations of the simulation framework

A critical issue in interpreting the absence of permeation in WT hERG, is whether it reflects the relatively low conductance of the channel and the insufficient sampling accessible. Under high-K^+^ conditions hERG displays a single-channel conductance of approximately 10 pS (Kiehn et al. 1996; Tan et al. 2022), corresponding to an ionic current of approximately 2 pA at -200 mV and to the translocation of roughly 12-13 K^+^ ions per microsecond. Although direct comparison between simulations and experiments must consider that the conventional additive force fields typically underestimate ion conductance due to the absence of explicit electronic polarization (Matamoros et al. 2023; Li et al. 2021; Hui et al. 2024), even assuming a conservative tenfold reduction of permeation rates would still predict detectable conduction events within the cumulative simulation time (20 μs per condition). Therefore, the complete absence of sustained permeation in WT hERG is unlikely to arise from insufficient sampling and instead reflects the intrinsic properties of the conformational ensemble adopted by its SF.

To further exclude limitations associated with the electrostatic description of the force-field, we repeated the simulations using the ECC framework, which partially accounts for electronic polarization and has been shown to improve agreement between simulated and experimental conductance in K^+^ channels (Vijayvergiya et al., 2015; Hui et al., 2025). Importantly, ECC substantially enhanced ion permeation in the N629D mutant, demonstrating that the simulation protocol can capture conduction when the filter adopts an appropriate architecture. In contrast, WT hERG remained completely non-conductive despite the improved electrostatic treatment. Thus, the ability of ECC to increase conduction in N629D but not in WT strongly argues that the absence of conduction is not a consequence of insufficient sampling or force-field limitations but rather arises from the structural properties of the WT SF ensemble.

### WT hERG spontaneously evolves toward a non-conductive inactivated-like SF conformation

A central finding of this work is that structural compatibility with ion coordination does not necessarily translate into a dynamically stable conductive state. Although the high-K^+^ WT hERG cryo-EM structures display a SF geometry resembling that of conductive K^+^ channels, this arrangement is intrinsically unstable when allowed to evolve without restraints, consistent with previous computational observations (Pettini et al., 2023). Across all simulated conditions, the SF reproducibly relaxed toward a structurally distinct conformation characterized by extracellular dilation around F627, localized constriction near V625, depletion of outer ion-binding sites, and persistent trapping of K^+^ ions within the central S2-S3 region.

These structural rearrangements provide a mechanistic explanation for the loss of conduction. Outward rotation of the F627 carbonyl groups disrupts the canonical ion-coordination geometry of the extracellular binding sites, leading to depletion of K^+^ ions from the S0-S1 region. Similar conformational changes have been observed in previous simulations of TREK two-pore-domain K^+^ channels, where rearrangement of the corresponding phenylalanine residue within the SF was identified as an initial conformational event associated with filter gating (Türkaydin et al. 2024). Simultaneously, constriction near V625 generates an additional barrier to ion translocation, as also shown in a recent study in which it is reported that wider SF allows higher conduction in MD simulations (Liu et al. 2026). Together, these changes disrupt the multi-ion occupancy pattern required for efficient knock-on conduction and promote persistent ion trapping within the central binding sites. Thus, we propose that conduction is abolished not by complete pore collapse but rather through remodeling of the SF into a configuration that can no longer sustain the coordinated ion dynamics required for K⁺ permeation.

### A possible structural correlate of early hERG inactivation

The non-conductive SF conformation identified here provides structural framework for interpreting hERG inactivation. The pronounced extracellular dilation closely resembles the expanded SF architecture recently resolved in inactivated Shaker channels, where loss of occupancy at the outer binding sites disrupts the conductive ion-coordination network (Stix et al., 2023). Unlike Shaker, however, WT hERG also develops a localized constriction near V625, producing a SF architecture that combines features of the dilated Shaker filter and the collapsed KcsA filter. Rather than conforming to either established paradigm, WT hERG therefore appears to sample a distinct intermediate within the structural landscape of C-type inactivation.

This interpretation is consistent with electrophysiological studies indicating hERG inactivation proceeds through multiple conformational states with distinct permeation properties. In the absence of extracellular K^+^, inactivated hERG channels transiently conduct Na^+^ before evolving into a more stable non-conductive state (Gang e Zhang 2006), suggesting that disruption of K⁺ conduction may precede complete loss of pore permeability. Although Na^+^ permeation was not examined here, the WT hERG SF conformation identified in our simulations provides a plausible explanation for an early inactivated intermediate. Specifically, the SF remains physically open but no longer supports the coordinated ion occupancy required for efficient K^+^ knock-on conduction.

Within this framework, the conformation identified here may correspond to, or at least share key structural features with, the early P-type-like inactivated state inferred from electrophysiological studies. The more extensive SF rearrangements recently observed in low-K^+^ cryo-EM structures, including disruption of central ion-binding sites and stabilization of alternative backbone conformations (Lau et al., 2024), may therefore represent later and more stable stages of the same inactivation pathway.

### N629D stabilizes the conductive SF state

A particularly strong line of evidence linking the observed SF distortion to inactivation comes from the behaviour of the N629D mutant. This mutation has long been known experimentally to strongly suppress fast inactivation, yet the structural basis of its effect has remained incompletely understood. In our simulations, N629D prevents the SF rearrangements observed in WT hERG, maintains a conductive filter architecture, and sustains robust voltage-dependent K^+^ permeation.

Unlike the WT hERG, the mutant maintains a stable SF architecture at both the extracellular F627 region and the central V625 region, that closely overlaps with those of conductive K^+^ channels such as KcsA, preserves occupancy of the outer ion-binding sites and supports continuous ion exchange across the SF. Importantly, the conductive architecture remains stable under both conventional and ECC conditions, whereas ECC selectively enhances permeation without altering SF structure, confirming that the increased ion transport reflects an intrinsically conductive SF rather than an artifact of the electrostatic model. Together, the suppression of inactivation, preservation of a canonical conductive SF architecture, and recovery of robust K^+^ permeation provide strong mechanistic supports for the interpretation that the conformational changes identified in the WT hERG are linked to the inactivation process. Rather than directly promoting conduction, N629D appears to stabilize the conductive SF ensemble and prevent progression toward the non-conductive intermediate sampled by WT hERG.

### Limitations and conclusions

Although the observed behaviour is highly reproducible across membrane potentials, and electrostatic treatments, MD simulations remain limited by accessible timescales and may not fully capture slower transitions between distinct inactivated conformations. Accordingly, the SF architecture identified here should not be viewed as the sole structural manifestation of hERG inactivation, but rather as one intermediate within a broader conformational landscape that may also include the more extensively disrupted SF states observed experimentally under low-K^+^ conditions. Future experimental studies will ultimately be required to establish the precise relationship between these structural intermediates and the functional states defined by electrophysiology.

Despite these limitations, our results challenge the prevailing view that the currently available high-K^+^ WT hERG cryo-EM structure represents a stable conductive state. Instead, our simulations indicate that this conformation spontaneously relaxes into an inactivated-like SF architecture incapable of supporting K^+^ permeation.

## Funding Sources

Funded by the European Union - NextGenerationEU under the National Recovery and Resilience Plan (PNRR) - Mission 4 Education and Research - Component 2 From Research to Business-Investment 1.1, Notice Prin 2022 - (DD N. 104 del 2/2/2022) title ‘‘Kinetic models of ion channels: from atomic structures to membrane currents’’, proposal code 20223XZ5ER - CUP J53D23006940006

## Acknowledgments

We acknowledge CINECA for awarding access to computational resources through the ISCRA Initiative (grant numbers HP10B597KB and HP10B5IPGG).

## Notes

### Competing Interest Statement

The authors have declared no competing interest.

