## Supporting Information for "The wild-type hERG cryo-EM structure fails to support conduction and evolve toward an inactivated-like selectivity filter conformation in molecular dynamics simulations"

| Model | Force Field | Membrane Potential | Simulations | Conduction events |
|---|---|---|---|---|
| hERG | Amber14sb | 200 mV | 4 x 5 µs | 0 |
| hERG | Amber14sb + ECC | 200 mV | 4 x 5 µs | 0 |
| hERG-N629D | Amber14sb | 200 mV | 4 x 5 µs | 2 partial |
| hERG-N629D | Amber14sb + ECC | 200 mV | 4 x 2 µs | 1 + 6 partial |
| hERG | Amber14sb | -200 mV | 4 x 5 µs | 0 |
| hERG | Amber14sb + ECC | -200 mV | 4 x 5 µs | 0 |
| hERG-N629D | Amber14sb | -200 mV | 4 x 5 µs | 1 partial |
| hERG-N629D | Amber14sb + ECC | -200 mV | 4 x 2 µs | 6 partial |
| hERG | Amber14sb | 400 mV | 4 x 5 µs | 0 |
| hERG | Amber14sb + ECC | 400 mV | 4 x 5 µs | 0 |
| hERG * | Amber14sb | 400 mV | 4 x 5 µs | 1 partial |
| hERG-N629D | Amber14sb | 400 mV | 4 x 5 µs | 2 + 2 partial |
| hERG-N629D | Amber14sb + ECC | 400 mV | 4 x 2 µs | 9 |
| hERG | Amber14sb | -400 mV | 4 x 5 µs | 1 partial |
| hERG | Amber14sb + ECC | -400 mV | 4 x 5 µs | 2 § |
| hERG-N629D | Amber14sb | -400 mV | 4 x 5 µs | 33 |
| hERG-N629D | Amber14sb + ECC | -400 mV | 4 x 2 µs | 24 |

**Table S1. Molecular Dynamics Simulations.** §The conduction events observed in simulations at -400 mV using the ECC parameters are not coherent with a sustained conduction mechanism (details in Figure S1).

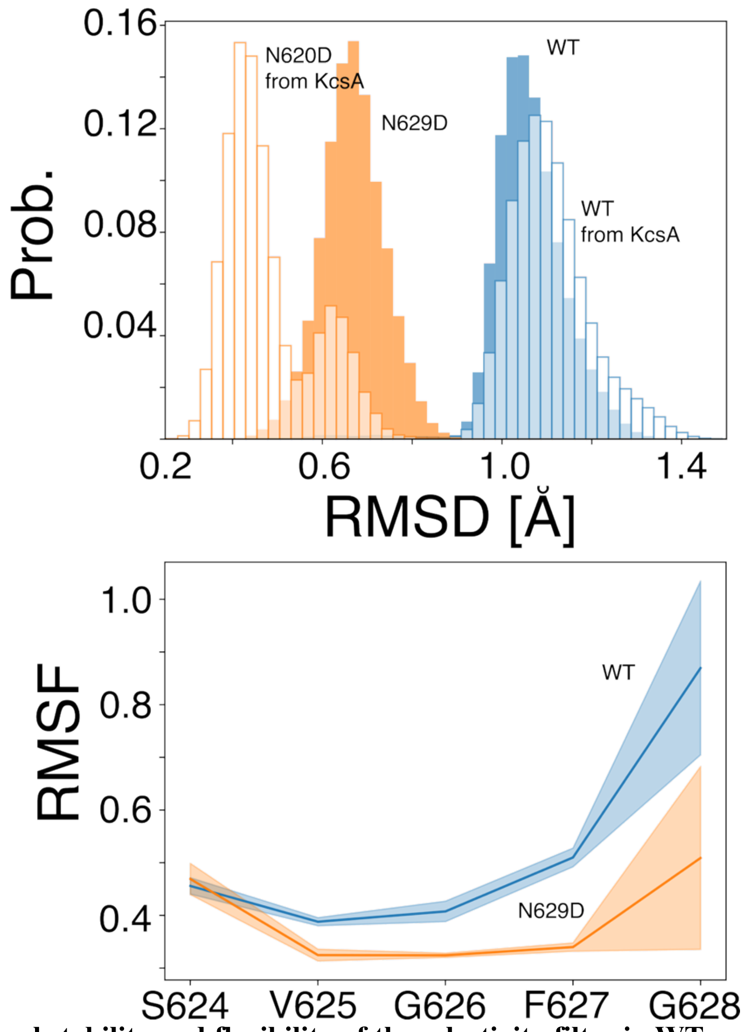

**Figure S1. Structural stability and flexibility of the selectivity filter in WT and N629D hERG.** (Upper panel) Probability distributions of the RMSD of the SF backbone atoms. Filled boxes represent the RMSD relative to cryo-EM structure (PDB ID: 9CHP), whereas empty boxes represent the RMSD relative to the X-ray structure of the KcsA channel in the conductive state (PDB ID: 1K4C). (Lower panel) RMSF of the alpha-carbon atoms of SF residues. Data were calculated from the four independent replicas simulated at +200 mV with Amber14SB force field and ECC parameters.

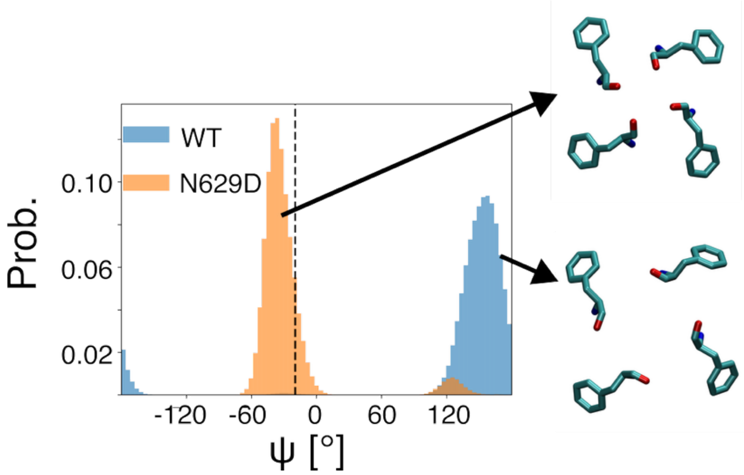

**Figure S2. Probability distribution of the backbone ψ -angles of residues F627.** The dashed horizontal line indicates the value observed in the cryo-EM structure. Representative configurations of the F627 residues, viewed from the extracellular side of the channel, are shown in licorice representation. The F627 carbonyl oxygens are oriented towards the pore axis in the N629D mutant but point laterally in the WT channel. Probability distributions were calculated from the four independent replicas simulated at +200 mV with Amber14SB force field and ECC parameters.

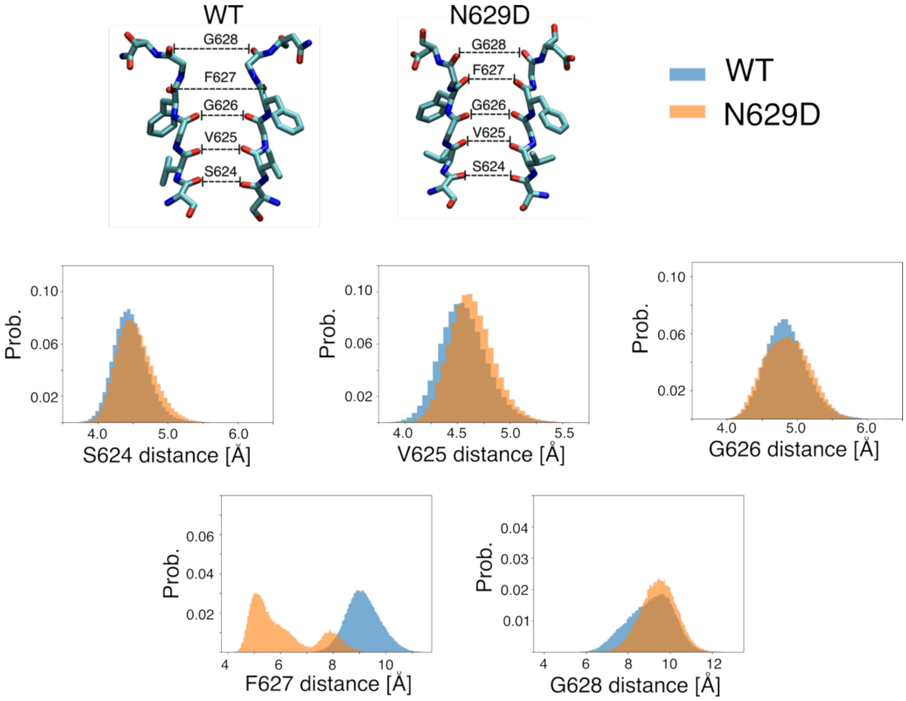

**Figure S3. SF architecture of WT and N629D hERG.** Probability distributions of the distances between carbonyl oxygen atoms of equivalent residues (S624, V625, G626, F627, and G628) from two opposing subunits are for WT hERG (blue), N629D hERG (orange). Data were obtained from simulations performed at +200 mV for all channels using Amber14SB force field and ECC parameters. The distance between opposing F627 carbonyl oxygen atoms is greater in WT than in N629D, consistent with simulations performed using Amber14SB without ECC parameters (Figure 4). The distribution of distances between opposing V625 carbonyl oxygen atoms is slightly shifted toward lower values in N629D relative to WT. This effect was more pronounced in simulations without ECC parameters, which also showed a similar decrease for G626. These differences may arise from the weaker electrostatic attraction between potassium ions and oxygen atoms when ECC parameters are used.

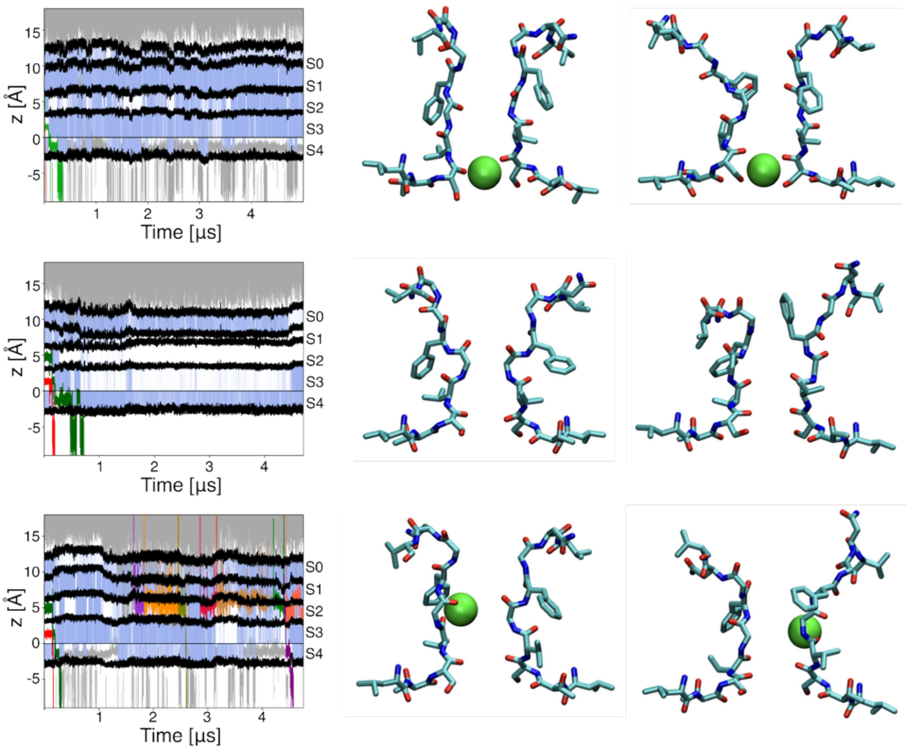

**Figure S4. Structural changes in simulations of the wild-type channel with ECC parameters at membrane potential equal to -400 mV.** The position of $K^+$ ions along the pore axis of the channel is shown using grey lines for ions outside the SF, and lines of different colors for ions inside the SF. Blue shading indicates binding sites, S0-S4, occupied by water molecules. Black lines correspond to the boundaries between binding sites of the SF. On the right side, residues 622 to 630 for the two pairs of opposing subunits are shown in licorice representation, with $K^+$ ions as green VdW spheres.
